# Simultaneous zero echo time fMRI of rat brain and spinal cord

**DOI:** 10.1101/2025.03.20.644420

**Authors:** Hanne Laakso, Lin Wu, Sara Ponticorvo, Raimo A. Salo, Jaakko Paasonen, Ekaterina Paasonen, Mikko Kettunen, Russell L. Lagore, Lance DeLabarre, Ethan Polcyn, Gregor Adriany, Javier Istúriz, Dee M. Koski, Djaudat Idiyatullin, Olli Gröhn, Silvia Mangia, Shalom Michaeli

## Abstract

**Purpose:** Functional assessments of the central nervous system (CNS) are essential for many areas of research. Functional MRI (fMRI) typically targets either the brain or the spinal cord, but usually not both, due to the obstacles associated with simultaneous image acquisitions from distant fields of view (FOVs) with conventional MRI. In this work, we establish a novel MRI approach that enables artefact-free, quiet, simultaneous fMRI of both brain and spinal cord, avoiding the need for dynamic shimming procedures.

**Methods:** We utilized zero echo time (TE) Multi-Band-SWeep Imaging with Fourier Transformation (MB-SWIFT) technique at 9.4T in a simultaneous dual-FOV configuration and two separate radio frequency (RF) transmit-receive surface coils. The first coil covered the rat brain, while the second was positioned approximately at the T13-L1 level of the rat”s spinal cord with copper shielding to minimize the coupling between the RF coils. Eight Sprague–Dawley rats were used for hindlimb stimulation fMRI studies.

**Results:** Robust and specific activations were detected in both the brain and spinal cord during hind paw stimulation at individual and group levels. The results established the feasibility of the novel approach for simultaneous functional assessment of the lumbar spinal cord and brain in rats.

**Conclusion:** This study demonstrated the feasibility of a novel dual-FOV fMRI approach based on zero-TE MB-SWIFT and set the stage for translation to humans. The methodology enables comprehensive functional CNS evaluations of great value in different conditions such as pain, spinal cord injury, neurodegenerative diseases, and aging.

## Introduction

Functional magnetic resonance imaging (fMRI) has revolutionized the field of neuroscience by enabling the non-invasive visualization of hemodynamic and metabolic processes linked to neuronal activity *in vivo*. It plays a crucial role in mapping brain function and network connectivity in both humans and animals. With the rapid development of advanced MRI techniques, it has become feasible to evaluate microstructure and function with high spatial sensitivity and specificity. However, all currently available pulse sequences used in functional or structural MRI research have predominantly focused on one organ, such as the brain or the spinal cord. In recent years, there has been increasing interest in expanding the capabilities of fMRI to encompass the spinal cord.^1,2^ Despite the growing interest and advancements in acquisition and processing strategies,^3^ spinal cord fMRI remains underdeveloped compared to brain fMRI. Susceptibility artifacts arising from tissue characteristics, motion of the spinal cord and cerebrospinal fluid induced by cardiac and respiratory cycles, and the relatively small cross-sectional dimension of the spinal cord compromise the quality of the images.^3,4^

Only a limited number of fMRI studies combining brain and spinal cord fMRI have been reported^5– 12^, mostly including only midbrain regions and the cervical spinal cord. However, neither more extensive coverage of the brain and spinal cord nor simultaneous fMRI has been reported so far. In fact, despite the potential impact of a comprehensive fMRI approach for studying the central nervous system (CNS), achieving such a goal poses considerable challenges. These are largely attributed to technical obstacles with conventional MRI techniques, particularly the necessity for high magnetic field homogeneity across a large field of view (FOV) or several FOVs capable of encompassing both the brain and spinal cord simultaneously. To overcome this issue, per-slice dynamic shimming approaches have been proposed.^4,13^ However, they present drawbacks, including a notable extension of scanning time, limitations imposed by the settling-time of eddy currents, and applications primarily restricted to covering the cervical but not lower regions of the spinal cord. Furthermore, within a single repetition time, only sequential rather than interleaved FOV acquisitions of the brain and spinal cord have been achievable, hindering the realization of simultaneous acquisitions optimal for comprehensive connectivity analyses of the CNS.

In this work, we utilized zero echo time (zero-TE) MRI technique called Multi-Band-SWeep Imaging with Fourier Transformation (MB-SWIFT)^14,15^ modified to acquire signals from two FOVs (dual-FOV) simultaneously. MB-SWIFT, as other zero TE sequences, is minimally affected by field inhomogeneities, thus inherently addressing most of the above-mentioned issues due to high bandwidth. We previously demonstrated the feasibility and numerous technical benefits of MB-SWIFT for studying the function of the rat brain^16–19^ and spinal cord^20^ separately. We have also shown that the functional contrast in MB-SWIFT fMRI originates from the inflow of unsaturated blood^17,21^, differentiating it from the traditional blood oxygen level dependent (BOLD) contrast.^22^ Here we utilized this novel MRI approach for artefact-free, quiet fMRI simultaneously from the brain and lumbar spinal cord during hind paw stimulation, avoiding the need for dynamic shimming approaches.^13^

## Methods

### MB-SWIFT pulse sequence with dual FOV capabilities

In Figure 1, the schematic representation of the radial pulse sequence modified from the original MB-SWIFT technique for dual-FOV acquisitions is presented. The pulse sequence allows for virtually simultaneous (up to ∼1 ms repetition time per spoke) acquisitions of two independent FOVs. As in the original MB-SWIFT method, the orientation of the gradient changes incrementally from spoke to spoke. During a given gradient orientation, sequential excitations and readouts are performed for the independent FOVs irradiated by two different RF coils. Acquisition is then repeated N_S_times, and thus the dual-FOV volume is acquired with an overall N_S_T_R_time resolution, where T_R_ is the repetition time between the excitation of the same FOV (i.e. T_R_ = 2 x spoke repetition time). The offset frequencies of the transmitters change according to the location of FOV. The amplitude of the readout gradient can be separately adjusted to allow the different sized FOVs. The number of pulses (number of gaps in the gapped pulse) can be increased to gain SNR and power efficiency or decreased for faster imaging. The MB-SWIFT with the dual-FOV capabilities is freely available for non-commercial use at https://license.umn.edu/product/swift-software-for-bruker-mri-systems#! for Bruker and at https://license.umn.edu/product/vmnrj-swift-software-for-agilent-varian-systems#! for Agilent platforms, respectively.

**Figure 1.**
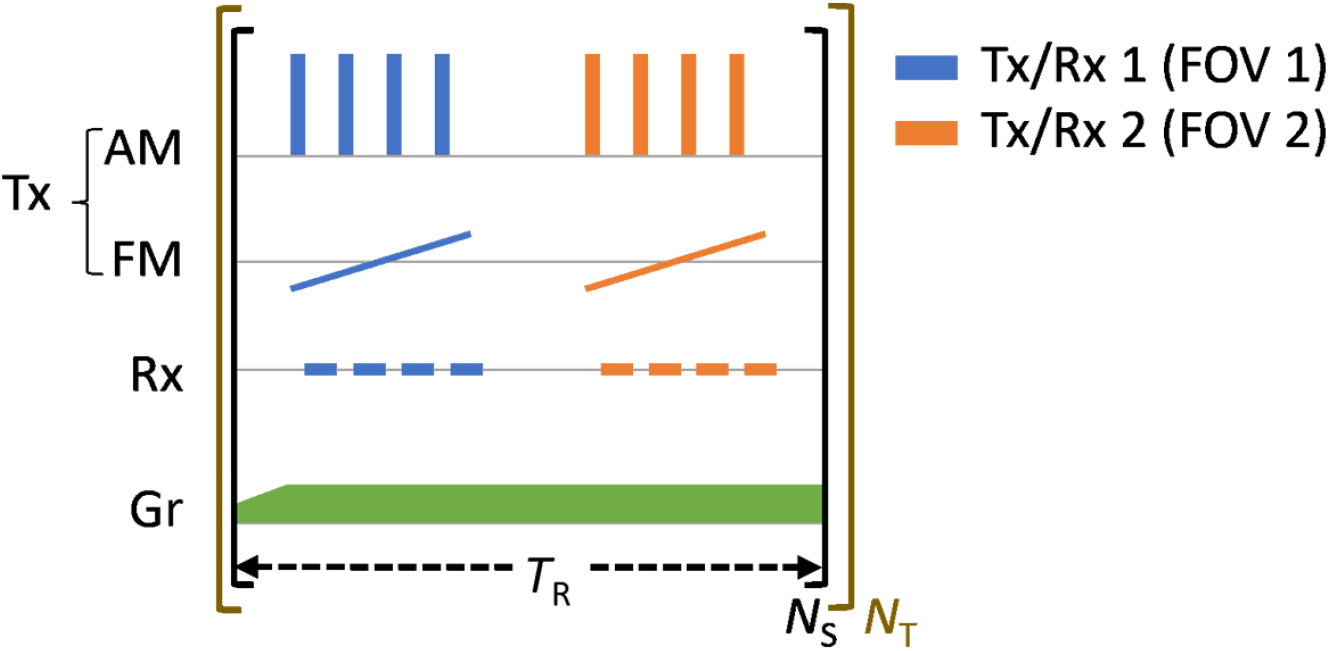
Schematic view of the dual FOV MB-SWIFT sequence for simultaneous acquisition of the two FOVs. T_R_: repetition time for the two spokes, N_S_: number of spokes to acquire one image volume, N_T_: number of time points (image series), AM: amplitude modulation function, FM: frequency modulation function, Tx1 and Tx2 transmitting channels, Rx1 and Rx2 receiving channels, Gr: field gradients, a change in the gradient amplitude between the spokes is required if different FOV sizes are used.

### Hardware configuration for MB-SWIFT fMRI

Figure 2A shows the setup and connections for the dual-FOV imaging at our scanner. Two RF amplifiers are required to be alternated accordingly to transmit the signal to a given coil, and for this, a Dual Amplifier Blanking Selection Unit (DABSU) was developed because our system does not support parallel transmission (pTx). The DABSU enables alternately unblanking two independent RF amplifiers for excitation two distant FOVs (Figure 2). In the instruments where pTx is available, transition of the signal to two separate coils could be achieved without DABSU.

**Figure 2.**
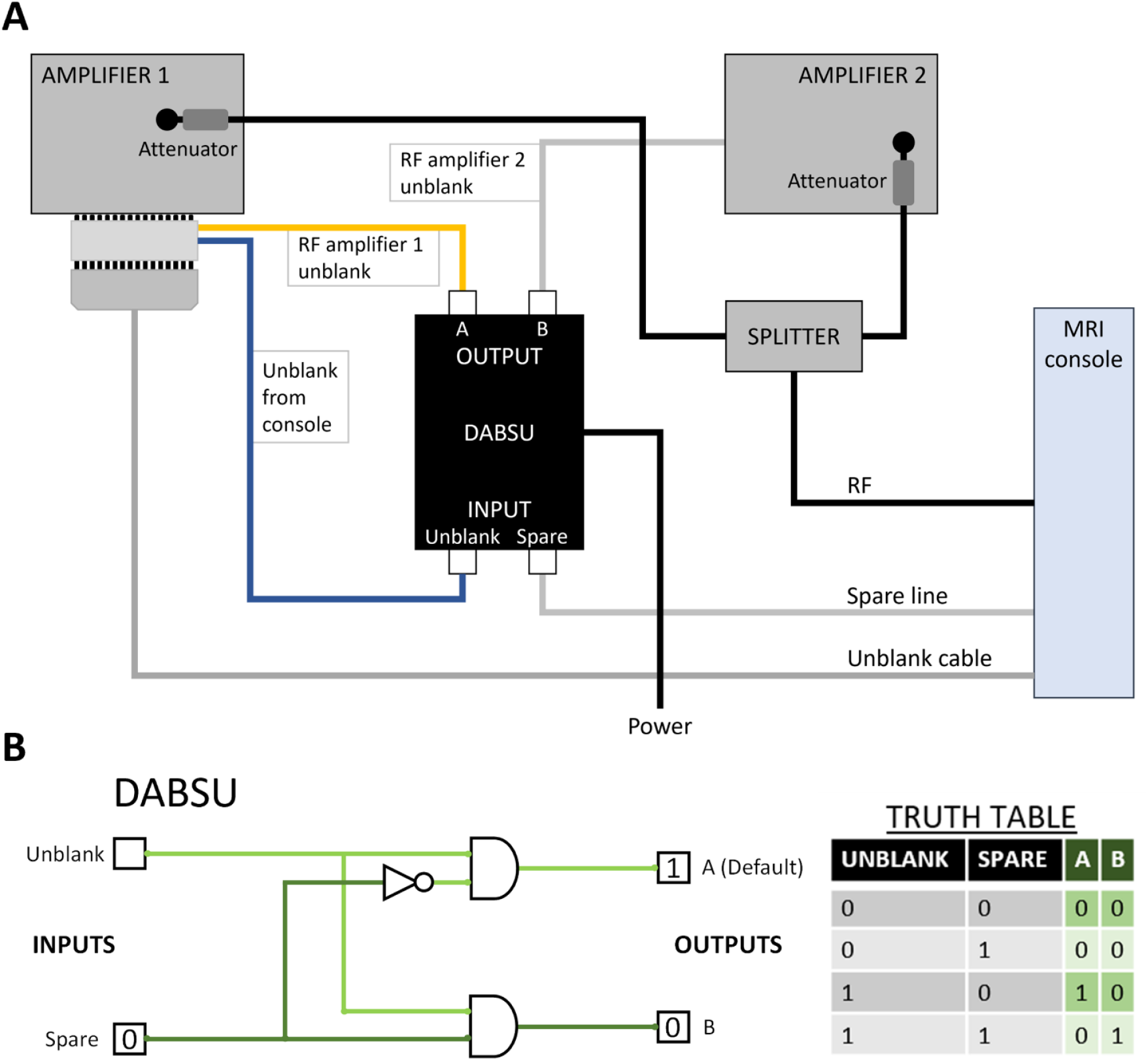
Schematic overview of the setup for dual-FOV. A) Connections from the console to Dual Amplifier Blanking Selection Unit (DABSU) controlling the unblanking of the two individual amplifiers. B) Schematic diagram of Dual Amplifier Blanking Selector Unit (DABSU). The input “Unblank” signal is distributed to output channel “A” or “B” depending on the signal level on input called “Spare”. The “Unblank” and “Spare” signals are supplied from the scanner”s console. The “A” and “B” outputs are connected to the “Unblank” inputs of two amplifiers.

The DABSU unit is a simple 5-volt digital logic device based on 7400-series (Texas Instruments) integrated circuits which intercepts the transistor-transistor logic (TTL) unblank signal from the console and also accepts an additional (spare) TTL signal from the console, and uses these inputs to generate two alternating unblank signals for the two independent RF amplifiers (Figure 2B). This enables alternated independent excitation of two FOVs. The truth table presented in Figure 2 shows the two outputs (A and B) produced from the two inputs (spare and unblank). Unblank goes high during the transmit pulse, while the spare is switched either low to unblank the RF amplifier connected to output A, or high to unblank the RF amplifier connected to output B.

Two individual transmit-receive RF loop coils were used at 9.4T equipped with the Agilent DirectDRIVE console (Palo Alto, CA, USA). The first oval shaped coil (inner axes 14 x 16 mm, Neos Biotec, Pamplona, Spain) covered the rat brain, while the second coil (inner axes 14 x 18 mm, Neos Biotec, Pamplona, Spain) was positioned approximately at the T13-L1 vertebral level of the rat spine with an adjustable RF shield loop (inner axes 34 x 62 mm, copper wire) placed around it to minimize coupling between the coils (Figure 3). This shielding loop consists of a closed loop of copper wire, only interrupted by a DC isolation capacitor, to prevent the flow of gradient-induced eddy currents, while allowing a free circulation of radio-frequency currents. The position of this RF shield loop is adjusted to an optimum spot where the magnetic flux directly coupled between head and spine coils cancels out with the flux indirectly coupled between them (through the circulation of an induced current in the shielding loop). Using the RF shield, we were able to achieve average decoupling of ∼-40 dB as measured on a network analyzer (NanoVNA-F V2, SYSJOINT Information Technology Co., Ltd., Hangzhou City, Zhejiang Province, China).

**Figure 3.**
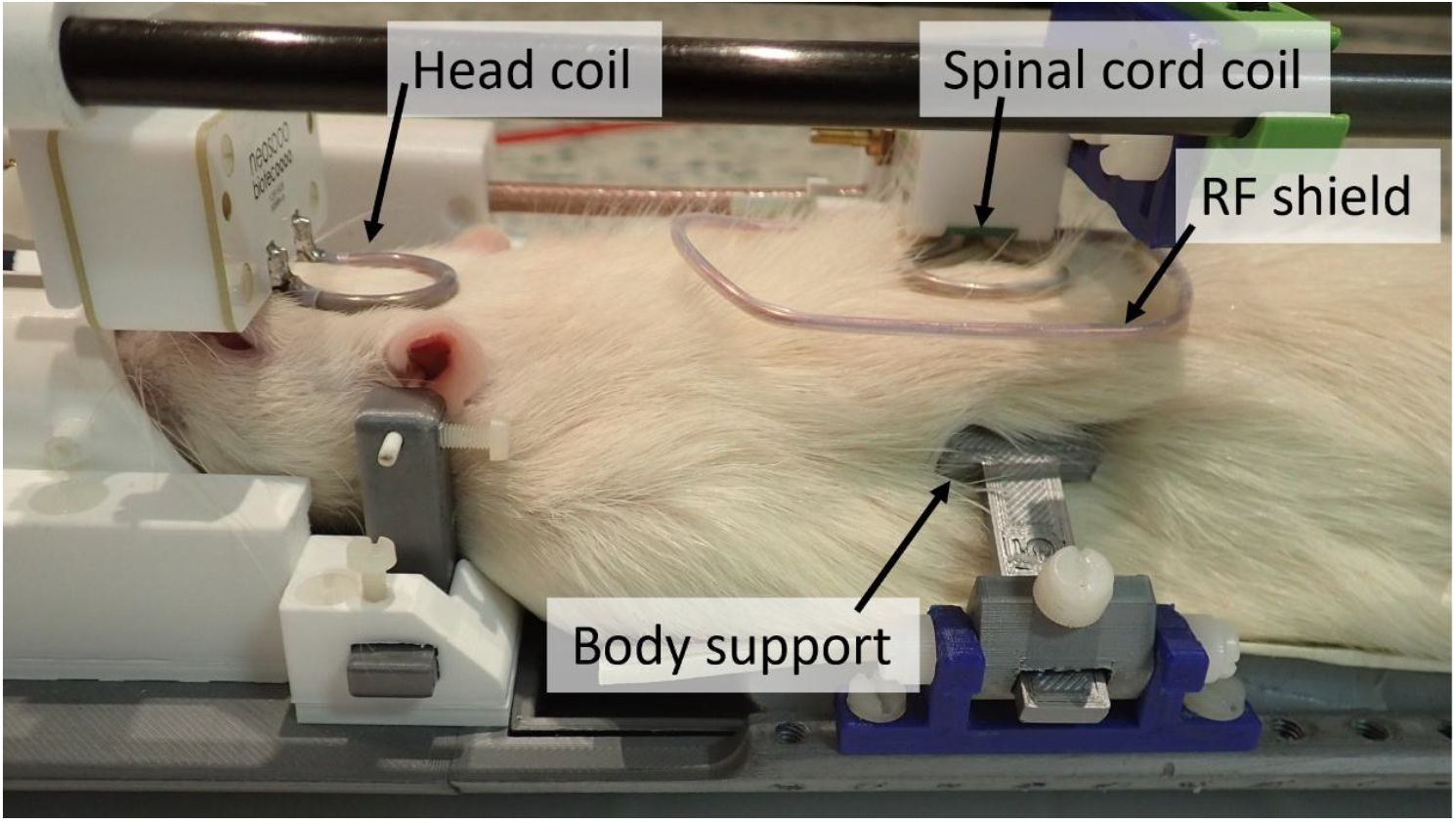
Setup for simultaneous brain and spinal cord fMRI. One oval shaped loop coil (inner axes 14 x 16 mm) was placed on the head and another oval shaped loop coil (inner axes 14 x 18 mm) was placed on the spinal cord of a rat at the vertebral level T13-L1. An adjustable RF shield loop (inner axes 34 x 62 mm, copper wire) was placed around the spinal cord coil to minimize coupling between the two coils. The body of the rat was supported with bars stabilizing the spine from the sides to reduce the motion from breathing.

For dual-FOV acquisitions we performed first order shimming based on the linewidth of spectra from signals of each FOV. Mostly Z-shimming gradient was adjusted to reach the same central offset frequency for both channels. This was achieved by performing global shimming on a large slab covering both brain and spinal cord, referred here to as average shim.

### Animal experiments

Eight Sprague-Dawley rats (weight 305 g ± 49 g (mean ± SD), 3 females and 5 males) were used for the studies. Food and water were available ad libitum. All animal procedures were approved by the Finnish Animal Experiment Board and conducted in accordance with the European Commission Directive 2010/63/EU guidelines.

The rats were initially anesthetized with isoflurane (5% induction, 2% during setup, carrier gas O_2_/N_2_= 30/70%) and then a bolus of 0.015 mg/kg medetomidine was given subcutaneously. The animal was placed on the holder and the head was fixed with toothbar and earbars, while the body of the rat was supported with bars stabilizing the spine from the sides to reduce breathing-induced motion (Figure 3). A warm water circulating heating pad was placed under the animal to keep the animal”s temperature close to 37 °C. An infusion line with a needle was inserted subcutaneously into the back of the rat and after 15 minutes from the medetomidine bolus, an infusion of medetomidine, 0.03 mg/kg/h, was started and the isoflurane was reduced to about 0.5%. Two needle electrodes were placed under the skin on the sides of the right heel pointing upwards to the leg of the animal, to stimulate the fascicles of the sciatic nerve. Before scanning, the stimulation was tested to observe slight movement of the paw induced by the electrical stimulation. Respiration and temperature of the animal were monitored during the scans using a pneumatic pillow under the animal and a rectal temperature probe (Model 1025, Small Animal Instruments Inc., Stony Brook, NY, USA), respectively. After the experiment, atipamezole, 0.5 mg/ml, 0.5 mg/kg was given to reverse the sedative effect of medetomidine.

### fMRI

MB-SWIFT technique modified for simultaneous dual-FOV acquisitions was used for fMRI acquisitions with 3 s temporal resolution and 1547 spokes per image volume, acquired in simultaneous fashion (interleaved spokes) with spiral trajectory. The other parameters were T_R_ =1.94 ms (single spoke acquisition time was 0.97 ms), four RF gaps, excitation/acquisition bandwidth (BW) = 192/384 kHz, FOV = 40 x 40 x 40 mm^3^, flip angle = 6°, matrix = 64^3^, and 625 µm isotropic resolution for both brain and spinal cord. In total 248 volumes per FOV were collected during the fMRI scans, resulting in 12 min 24 s acquisition time.

An anatomical high-resolution image was acquired using magnetization transfer (MT) weighted MB-SWIFT with similar parameters, but with 4000 spokes per spiral, 16 spirals, 8° flip angle, and 256^3^matrix, 156 µm isotropic resolution. For MT, a *Sinc* pulse placed at 1500 Hz off-resonance was utilized. Pulse duration of the *Sinc* pulse was 15 ms, given every 32 spokes at the amplitude γB_1_ = 167 Hz. The total acquisition time for the anatomical scan was 8 minutes for the two FOVs.

In four rats, we also conducted comparative scans using single-FOV MB-SWIFT acquisitions with time resolution of 1.5 s and 496 volumes in the brain and spinal cord separately. For those acquisitions, the flip angle was reduced to 4° to compensate for the effect of shorter repetition time to achieve the same Ernst condition.

Furthermore, we performed an additional experiment in one animal with dual-FOV and single-FOV MB-SWIFT and comparative SE-EPI acquisitions with parameters TR = 3 s, TE = 35 ms, FOV = 40×40 mm^2^, 18 slices 1 mm thick. Two shimming strategies were used for EPI: first, averaged for both coils as in dual-FOV, and second, targeted for the imaged FOV.

The stimulation paradigm started with 60 s of rest followed by a 24 s block of stimulation and 90 s of rest repeated 6 times. The stimulation block consisted of 600 μs symmetric charge balanced pulses at 9 Hz frequency and 2 mA current. We used STG4008 stimulus generator (Multi Channel Systems MCS GmbH, Reutlingen, Germany) in the current mode to deliver the stimulation paradigm which was synchronized to start with the first volume of the fMRI acquisition by a TTL trigger pulse from the scanner.

### Reconstruction and preprocessing

The MB-SWIFT data were reconstructed as in ^23^ using RF-pulse deconvolution, gridding and iterative FISTA algorithm^24^ with 13 iterations. All MRI data were processed and analyzed with in-house made scripts, using Snakemake^25^ (https://snakemake.github.io/), Python (version 3.10, https://www.python.org/downloads/), advanced normalization tools^26^ (ANTs; https://stnava.github.io/ANTs/), FSL (version 6.0, https://fsl.fmrib.ox.ac.uk/fsl/fslwiki/), and FSL FEAT.^27^

Motion correction was performed by aligning the time series images to the first volume using ANTs.^28,29^ An independent component analysis (ICA; https://fsl.fmrib.ox.ac.uk/fsl/fslwiki/MELODIC) based motion regression was conducted to further improve the data quality by removing components representing motion, similarly as in ^30^. The motion components were automatically selected from the obtained 15 components and regressed out. Motion in the image time-series was estimated by the movement of the signal center of mass in three dimensions. To define the component as motion, two criteria were set: (1) when the correlation of the time series of the component with one of the motion parameters exceeded 0.75, and (2) when more than 70% of the spatial map voxels were located on the edge regions of the brain or outside, on top of the spinal cord.

Co-registration of the anatomical images for group level analysis was done in two steps using ANTs; First, separately for the brain and spinal cord images, five landmarks were selected and labeled from each of the images to make an initial rigid co-registration to reference brain and spinal cord selected from the data. Second, a nonlinear SyN registration was applied to refine the rigid registration. Subsequently, the registration transformations were applied to the functional data.

For the separate data set collected for comparison with EPI, motion correction was performed with ANTs similarly as with the other data, but no ICA based motion regression was conducted for either EPI or MB-SWIFT. The EPI data was slice time corrected and motion correction was done using the same protocol with ANTs as with the MB-SWIFT data.

For the subject-wise activation maps, the time series were high-pass filtered (0.01 Hz) and the autocorrelation was removed. The maps were calculated in FSL FEAT using a general linear model (GLM) for which the stimulation periods were modeled as boxcar functions and convolved with a gamma-variate impulse response function (IRF) assuming delay of the response 2.8 s and variation 1.4 s resembling the previously reported IRFs for rodents.^21^ To enable fair comparison between single-FOV and dual-FOV acquisitions, from single-FOV MB-SWIFT data every other volume was taken into account for calculating the activation map with the same time resolution of the dual-FOV MB-SWIFT dataset. The group-level activation maps were estimated using the FSL nonparametric permutation tool Randomise^31^ with the threshold-free cluster enhancement test statistic, with p-values corrected for family-wise error (FWE) rate and values of p < 0.05 considered as significant.

## Results

The dual-FOV MB-SWIFT acquisitions using average shimming on both FOVs provided good image quality and stimulation-based functional contrast in both FOVs. Robust and specific activations were detected in both the brain and spinal cord during hind paw stimulation in 7 out of 8 rats (Figure 4). Specifically, in the brain, the responses localized to the contralateral somatosensory and motor cortex and ipsilateral cerebellum (Figure 4A). In the spinal cord the activated area was at T13-L1 vertebral level (L4 spinal) ipsilateral to the stimulation and mostly in the dorsal horn (Figure 4B). The cortical cluster size was 18 -39 voxels across animals with cluster-averaged z-value between 4.3-5.8 (Supplementary Table 1). In the spinal cord, the cluster size was 42-166 voxels across animals with cluster-averaged z-values 4.3-6.8 (Supplementary Table 1). At group level, we detected clear responses in the spinal cord at L4 spinal level in the ipsilateral dorsal horn and in the brain in the contralateral somatosensory and motor cortex (Figure 5A,B). The time courses obtained from the areas depicted by the group level activation also show clear response to the stimulation both in the brain (∼0.5% signal change) and in the spinal cord (∼1% signal change) (Figure 5C,D). The responses in brain and spinal cord show different time courses; the response peaks around 9 s after the stimulation start in brain and after around 18 s in the spinal cord (Figure 5E,F). Furthermore, the return to baseline appears to take longer in spinal cord (∼45 s) compared to brain (∼33 s). Dual-FOV MB-SWIFT and single-FOV MB-SWIFT data showed similar activation patterns both in the brain and spinal cord (Figure 6A-B) and no significant differences were found between the activations (p > 0.05, FWE corrected, Randomise).

**Figure 4.**
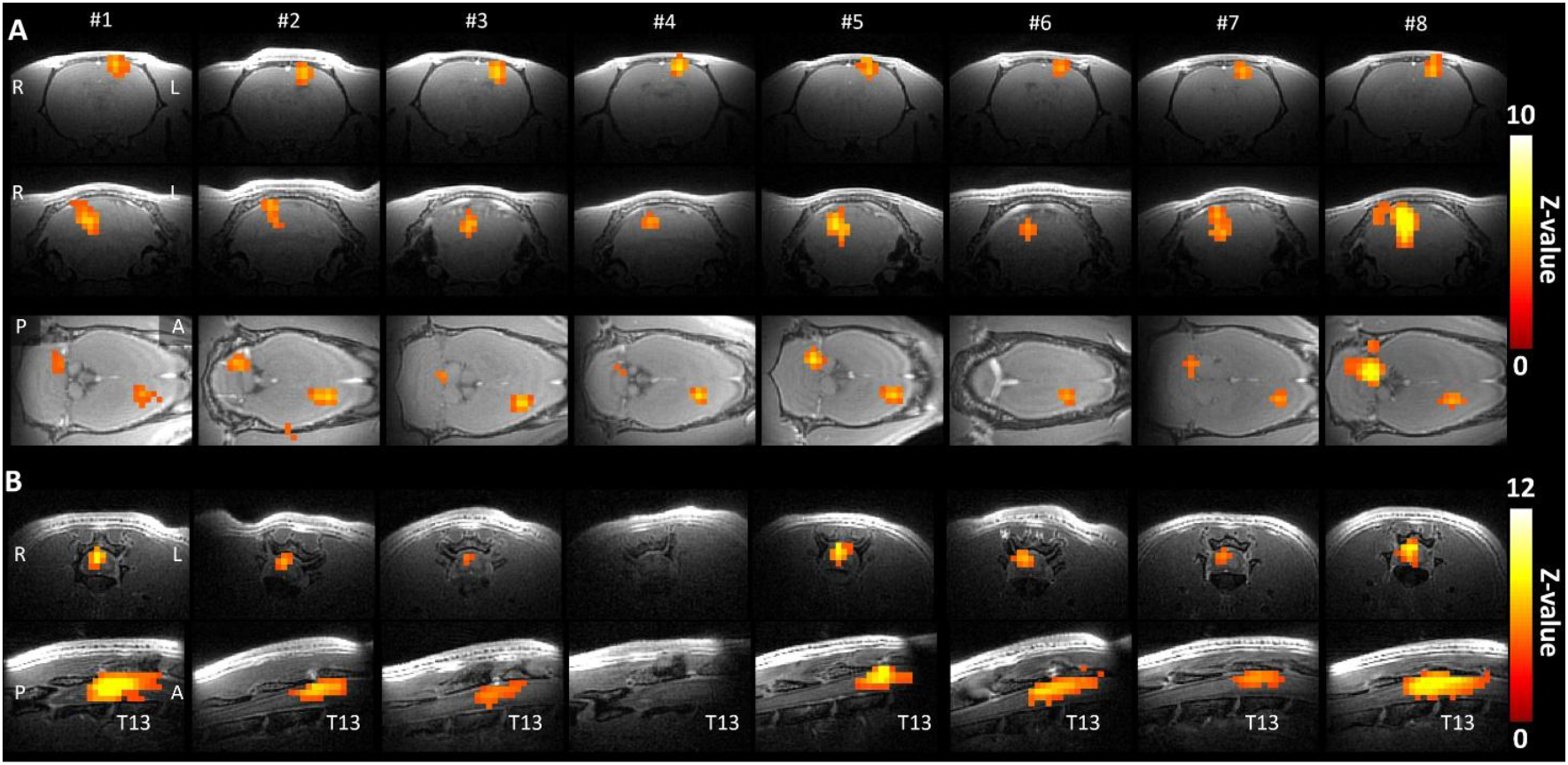
Dual-FOV MB-SWIFT fMRI activation maps from A) brain at cortex and cerebellum and B) spinal cord from individual animals during hind limb stimulation. In the brain, the activations are located in the contralateral somatosensory and motor cortex and ipsilateral cerebellum. In the spinal cord, the activation is ipsilateral, mostly on the dorsal side at T13-L1 vertebral level (L4 spinal). P<0.05, FWE-corrected. R=right, L=left, P=posterior, A=anterior, T13= the level of T13 vertebra.

**Figure 5.**
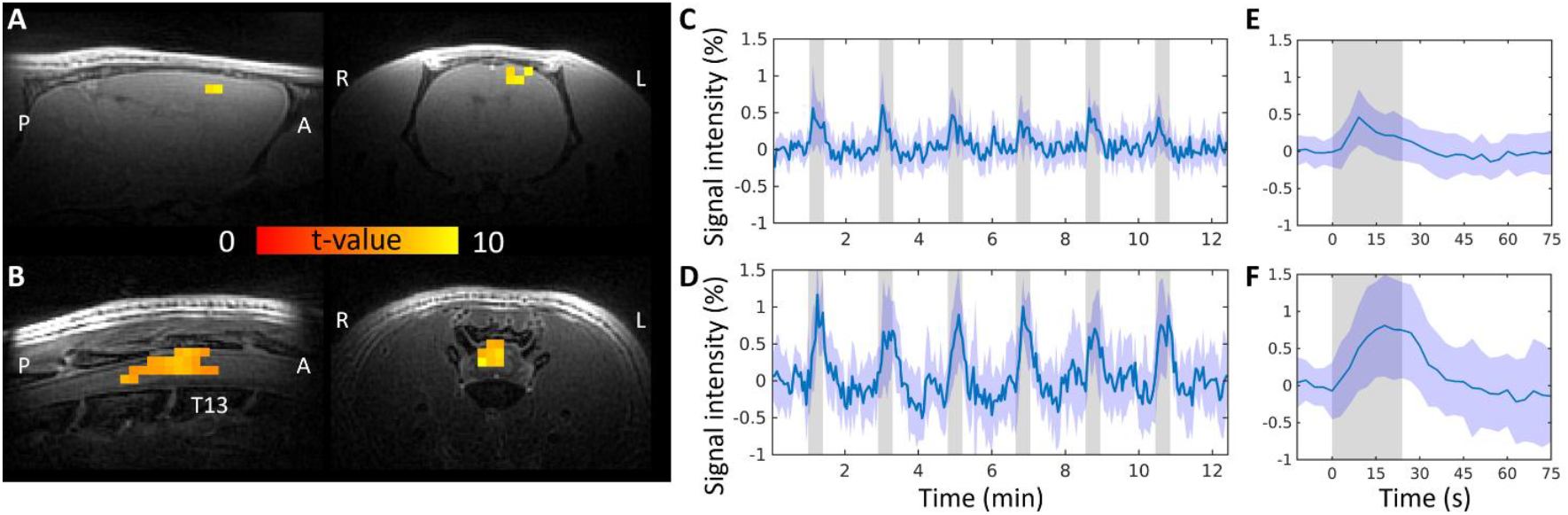
Group level analysis of dual-FOV MB-SWIFT fMRI during hind limb stimulation. The activation at group level is localized in the left somatosensory cortex (A) and right dorsal spinal cord at L4 spinal level (B). P<0.05, FWE corrected. The time courses (C,D) show the mean time courses from the activation clusters shown on the same row, and the shaded area represents the SD across subjects. The mean activation across subjects and scans for a single stimulation block is shown for brain (E) and spinal cord (F). The grey bars depict the stimulation timings. R=right, L=left, P=posterior, A=anterior, T13 = the level of T13 vertebra.

**Figure 6.**
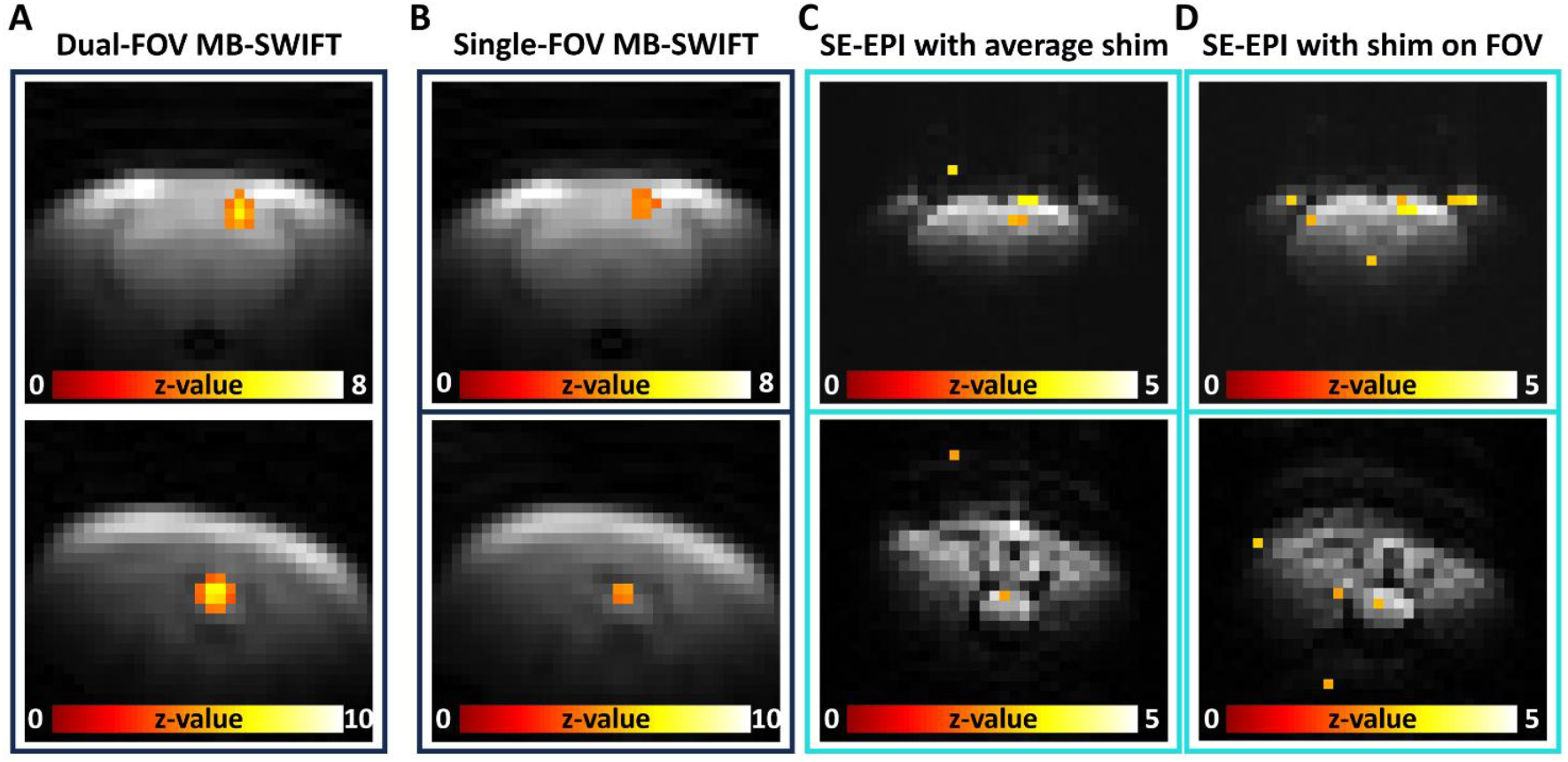
Comparison between dual-FOV and single-FOV MB-SWIFT and SE-EPI fMRI during hind limb stimulation. fMRI activation with: A) simultaneous MB-SWIFT acquisition from two FOVs with average shim; B) with separate single-FOV MB-SWIFT acquisitions with average shim; C) separate SE-EPI acquisitions with average shim; D) separate SE-EPI acquisitions with shimming performed on the target FOV. P<0.05, FWE corrected. The fMRI activation maps are overlaid on the original functional scans to show image quality.

When compared to EPI scans with average shim, MB-SWIFT images of the spinal cord were less distorted and enabled robust detection of activation in the spinal cord (Figure 6). On the other hand, only a few statistically significant voxels not forming a clear cluster were detected in the spinal cord with EPI and single activated voxels were seen elsewhere, also outside the rat body, indicating lower quality of data. When shimming individually to the imaged FOVs, i.e., brain or spinal cord, the EPI images were slightly less distorted than with average shim, but the activation patterns were still sparse as compared to MB-SWIFT (Figure 6).

## Discussion

In this study, we demonstrated that MB-SWIFT enables comprehensive stimulation-based fMRI studies of the CNS by acquiring signals virtually simultaneously from two distant sites, the brain and the lumbar spinal cord of a rat. The proposed approach involves the use of MB-SWIFT to excite two different FOVs using two independent coils and two RF amplifiers. Our data demonstrate the benefits of MB-SWIFT with dual-FOV capabilities compared to EPI for CNS fMRI without dedicated shimming approaches. Although we utilized the MB-SWIFT pulse sequence for simultaneous fMRI study of brain and spinal cord, the results of this work are generalizable also to other zero or ultra-short echo time MRI approaches. In general, the method could be used for simultaneous imaging of other organs as well.

MB-SWIFT offers the distinct benefit for imaging with virtually no echo time and high bandwidth thus inherently minimizing sensitivity to frequency offset variations. This is particularly beneficial for CNS fMRI because it enables circumventing the challenges of dynamic shimming required for large FOVs, and it minimizes sensitivity to both motion and signal dropouts due to susceptibility artefacts which are prominent in spinal cord imaging. Furthermore, with radial acquisitions, sampling of two FOVs can occur with a time-shift of a single spoke duration (i.e., within 1 ms of each other), and the center of k-space critical for detecting fMRI signals is consistently sampled in each spoke for each FOV. Overall, MB-SWIFT enables virtually simultaneous acquisition of two FOVs. Moreover, in MB-SWIFT, the gradients are on during the RF pulse which allows incremental gradient switching that minimizes gradient-induced artefacts during electrophysiological recordings and provides a nearly silent acquisition, enhancing subject comfort ^18^. Therefore, future applications of this method could be extended, for example, to MRI of different organs combined with multi-site electrophysiological recordings or to brain fMRI of different subjects positioned simultaneously in one magnet, i.e., social MRI. Such studies which are focused on brain fMRI of two mice in awake state are underway in our laboratory.

In this work, we detected activation patterns in both targets, the brain and spinal cord, in relevant areas in response to electrical hind limb stimulation. We stimulated the branches of sciatic nerve that originates from spinal cord levels L4 and L5,^32^ where we detected the spinal cord responses, supported also by previous findings.^33,34^ In accordance with previous brain fMRI studies during electrical stimulation of the hind limb, we observed functional activation in the somatosensory cortex^35–38^ and in cerebellum.^34,39^ However, we did not observe responses in thalamus, which is part of the sensorimotor network,^40^ activated especially with noxious stimulation.^41^ Lack of thalamic response is relatively common finding in studies in anesthetized animals with non-noxious sensory stimulus.^37,38^

The ability to detect an fMRI signal simultaneously from distinct sites along the sensory pathway is unique to this technique and could be particularly valuable, for example, for assessing recovery and treatment effects after spinal cord injury or in assessing pain processing. Additionally, when comparing the results obtained with dual-FOV *vs* single-FOV acquisitions, the functional contrast appears very similar, confirming that the quality of the images and the sensitivity to functional activity remains the same when acquiring multiple FOVs. Finally, when compared to the standard SE-EPI fMRI sequence, the benefit of MB-SWIFT is in obtaining artefact-free images with minimal shimming also from two FOVs simultaneously.

### Limitations

Several limitations of this study should also be indicated. For shimming, we paid particular attention to minimize the offset frequency for both channels simultaneously, considering that performing the shimming of an extended area covering both FOVs is very challenging. In our set-up, this choice was appropriate because the MB-SWIFT pulse sequence tolerates the line broadening if it is smaller than BW/N, where the BW is typically 192 kHz for MB-SWIFT, and N is the number of voxels along one dimension (frequently equal to 64 for fMRI experiments). So, the half maximum linewidth of the water proton signal must be smaller than 3 kHz, which is a very modest requirement, especially in comparison with EPI type acquisition which imposes a requirement for linewidths < 50-100 Hz.

Additionally, physiological noise and body movement reduce image SNR and fMRI contrast. We utilized spinal cord supports in our set-up to limit the effect of the breathing movement in the fMRI spinal cord signal, but further development of the set-up is needed to improve reproducibility of the spinal cord functional activity. With the current setup, we detected activation in the spinal cord in seven out of eight cases with statistical significance of p < 0.05, FWE corrected. Using a lower threshold, we could see activation also in that animal. The lack of significant activation was likely due to the movement (single larger movement, which was not removed by our motion correction protocol, Supplementary Table 2). The current analysis pipeline includes motion correction and removal of components representing motion, however other established approaches to model physiological noise in the fMRI data (e.g., RETROICOR^42^), could be explored to increase fMRI sensitivity in the spinal cord.

Finally, the size of the gradient coils in the Z direction relative to the object should also be considered. The maximal distance between FOVs is limited by the size of the X, Y and Z gradient coil of the system as well as the overall magnet homogeneity. If the location of the object is close to the edges of the gradient coil where the gradients start to decline, the images will be distorted. Due to the design of the gradient coils, the gradients decline faster in X and Y direction, and in some cases, this could lead to signal collapse to one bright point and create folding artifacts in the images, which can appear if the size of the rat is too large. To avoid this artifact, the RF excitation of the areas outside the primary

FOVs needs to be avoided or substantially minimized. However, with the current set-up, it is impossible to avoid this as the trunk of the animal will remain outside the spinal cord coil, meaning that there could be some signal coming from further away, where the gradients start declining. We dedicated particular attention to this aspect when positioning the FOVs. However, in some cases, the distortion of the image or the presence of one bright artifact point can be observed at the edge of the image, especially for rats of bigger dimension. A potential solution that needs to be explored in future studies could be based on software solutions allowing the artifact removal in postprocessing or improvement of the hardware by introducing additional shielding of the RF coils to limit the extent of the B1 field beyond the z-gradient.

## Conclusion

We demonstrated the feasibility of dual-FOV fMRI based on the zero-TE MB-SWIFT technique in task-based fMRI. This methodology opens a new investigational window to study the functional synchronization of brain and spinal cord activity during specific tasks. The method could be used for fMRI investigations of the CNS in basic and clinical neuroscience, including several disorders that manifest in both the brain and spinal cord such as chronic pain, spinal cord injury, and neurodegenerative diseases.

## Acknowledgements

This work was supported by the Research Council of Finland grants 331955 (HL) and Flagship of Advanced Mathematics for Sensing Imaging and Modelling 358944, NIH grants R01NS129739, P41 EB027061, S10 OD032192, and WM KECK Foundation. The work was carried out at Kuopio Biomedical Imaging Unit, University of Eastern Finland, Kuopio, Finland (part of Biocenter Kuopio, Finnish Biomedical Imaging Node, and EuroBioImaging). The computational analysis was performed on servers provided by UEF Bioinformatics Center and Biocenter Kuopio, University of Eastern Finland, Finland. We want to acknowledge Ing. Lenka Dvořáková for assistance in the coregistration of the spinal cord data.

