## Supplementary Table 1 for "Simultaneous zero echo time fMRI of rat brain and spinal cord"

**Supplementary Table 1. The cluster size and average z-value of the activated area in the cortex and spinal cord in individual animals.**

| Cortex | | | | | | | | |
|---|---|---|---|---|---|---|---|---|
| Animal # | 1 | 2 | 3 | 4 | 5 | 6 | 7 | 8 |
| Cluster size | 31 | 37 | 38 | 39 | 28 | 18 | 22 | 29 |
| Cluster-averaged Z-value ± SD | 4.6 ± 1.1 | 5.1 ± 1.5 | 5.0 ± 1.6 | 5.8 ± 1.9 | 4.7 ± 1.3 | 4.3 ± 0.9 | 4.7 ± 1.0 | 4.7 ± 1.2 |
| Spinal cord | | | | | | | | |
| Cluster size | 115 | 67 | 42 | - | 63 | 92 | 53 | 166 |
| Cluster averaged Z-value ± SD | 6.4 ± 2.9 | 5.4 ± 1.9 | 4.3 ± 0.9 | - | 5.7 ± 2.5 | 5.5 ± 1.8 | 4.5 ± 1.1 | 6.8 ± 3.4 |
