## Supplementary Table 2 for "Simultaneous zero echo time fMRI of rat brain and spinal cord"

**Supplementary Table 2. Parameters of relative motion in pixels in the spinal cord FOVs in individual animals before and after correcting for motion.**

| Spinal cord before motion correction | | | | | | | | |
|---|---|---|---|---|---|---|---|---|
| Animal # | 1 | 2 | 3 | 4 | 5 | 6 | 7 | 8 |
| Max relative movent in pixels | 0.133 | 0.040 | 0.180 | 0.206 | 0.170 | 0.098 | 0.281 | 0.191 |
| Mean relative movent ± SD in Z | 0.045 ± 0.028 | 0.020 ± 0.010 | 0.066 ± 0.055 | 0.048 ± 0.040 | 0.078 ± 0.042 | 0.060 ± 0.022 | 0.110 ± 0.088 | 0.076 ± 0.054 |
| Mean relative movent ± SD in Y | 0.004 ± 0.003 | 0.003 ± 0.003 | 0.003 ± 0.003 | 0.027 ± 0.019 | 0.009 ± 0.005 | 0.007 ± 0.004 | 0.007 ± 0.005 | 0.009 ± 0.006 |
| Mean relative movent ± SD in X | 0.011 ± 0.008 | 0.014 ± 0.008 | 0.044 ± 0.030 | 0.017 ± 0.011 | 0.027 ± 0.016 | 0.012 ± 0.009 | 0.024 ± 0.017 | 0.054 ± 0.028 |
| Spinal cord after motion correction | | | | | | | | |
| Max relative movent in pixels | 0.047 | 0.030 | 0.064 | 0.193 | 0.093 | 0.088 | 0.105 | 0.038 |
| Mean relative movent ± SD in Z | 0.012 ± 0.010 | 0.013 ± 0.006 | 0.026 ± 0.015 | 0.048 ± 0.046 | 0.030 ± 0.019 | 0.052 ± 0.023 | 0.041 ± 0.027 | 0.011 ± 0.007 |
| Mean relative movent ± SD in Y | 0.008 ± 0.006 | 0.004 ± 0.003 | 0.010 ± 0.006 | 0.013 ± 0.009 | 0.018 ± 0.008 | 0.006 ± 0.004 | 0.008 ± 0.004 | 0.005 ± 0.004 |
| Mean relative movent ± SD in X | 0.010 ± 0.005 | 0.005 ± 0.003 | 0.016 ± 0.007 | 0.013 ± 0.011 | 0.024 ± 0.013 | 0.013 ± 0.009 | 0.016 ± 0.012 | 0.006 ± 0.005 |
